# Minor allele frequency thresholds dramatically affect population structure inference with genomic datasets

**DOI:** 10.1101/188623

**Authors:** Ethan Linck, C.J. Battey

## Abstract

One common method of minimizing errors in large DNA sequence datasets is to drop variable sites with a minor allele frequency below some specified threshold. Though widespread, this procedure has the potential to alter downstream population genetic inferences and has received relatively little rigorous analysis. Here we use simulations and an empirical SNP dataset to demonstrate the impacts of minor allele frequency (MAF) thresholds on inference of population structure. We find that model-based inference of population structure is confounded when singletons are included in the alignment, and that both model-based and multivariate analyses infer less distinct clusters when more stringent MAF cutoffs are applied. We propose that this behavior is caused by the combination of a drop in the total size of the data matrix and by correlations between allele frequencies and mutational age. We recommend a set of best practices for applying MAF filters in studies seeking to describe population structure with genomic data.

## Introduction

The distribution of genetic variation within and among individuals is the crucial to understanding the organization of biological diversity and its underlying causes. Across the genome, the effects of different evolutionary processes and historical events can result in different classes of genetic variants characterized by their relative frequency in a given population. An excess of common alleles may reflect the signature of population bottlenecks (Marth et al. 2004), purifying selection (Fay et al. 2001), or the absence of population subdivision (Pritchard et al. 2000). Alternatively, high frequencies of rare alleles can provide evidence of population expansion (Marth et al. 2004), detailed information on mutation rates and gene flow (Slatkin 1985), and reveal geographically localized population subdivision (Barton and Slatkin 1986, Gombert et al. 2014). Because the distribution of allele frequencies across sites (also known as the site frequency spectrum, or SFS) reflects the unique combination of these varied factors, downstream analyses are sensitive to the influence of sampling methodologies that alter the SFS. Yet despite the explosive recent growth of population genetics provided by advent of affordable reduced-representation genome sequencing for nonmodel organisms, there remain significant gaps in our knowledge of how data collection biases population genetic inference.

These biases may originate either in wet lab or bioinformatic treatments. Prior to sequencing, the SFS may be shaped by ascertainment bias in library preparation: RADseq-style methods introduce genealogical biases (Arnold et al. 2013) and nonrandom patterns of missing data (Gautier et al. 2012) due to reliance on the presence of restriction cut sites, while hybridization capture with ultraconserved element (UCE) probesets necessarily involves targeting sites highly conserved across evolutionarily distant taxa (Faircloth et al. 2012). During sequencing itself, relatively high error rates are accepted in individual reads, under the assumption they will be corrected during bioinformatic processing steps (Nielsen et al. 2012). However, the absence of standard bioinformatic pipelines in ecology and evolutionary biology is itself a source of uncertainty (Shafer et al. 2016) because specific methodologies and parameter choices may dramatically affect the composition of data matrices.

For organisms lacking a suitable reference genome, *de novo* sequence assemblies may introduce substantial errors that affect both the SFS and inference of population genetic structure (Shafer et al. 2016). During read-mapping, SNP variation can result in higher rates of successful alignments in reads sharing the reference allele (Degner et al. 2009). Parameters used during variant detection can also play a significant role in determining the number and distribution of single nucleotide polymorphisms or SNPs (Neilsen et al. 2012), the most frequently used marker type in modern population genetics. In particular, minor allele frequency (MAF) thresholds directly influence the SFS by imposing a cutoff on the minimum allele frequency allowed to incorporate a specific genetic variant. But despite its potential importance, the two most popular comprehensive bioinformatic pipelines for RADseq data alternatively include (Catchen et al. 2013) or exclude (Eaton et al. 2014) the option to set minor allele frequency thresholds during variant calling, with the result that among empirical studies MAF thresholds are only sometimes reported (e.g., Winger 2017, Blanco-Bercial and Bucklin 2016).

One potential consequence of ambiguous MAF choice is variation in the ability to detect population subdivision (or structure), a fundamental goal of many population genetic studies. Broadly speaking, methods to detect population structure fall into two categories: model-based (or parametric) approaches, and nonparamatric approaches. Model-based methods, exemplified by the influential program structure (Pritchard et al. 2000), typically assume a hypothetical *K* populations characterized by *p* allele frequencies at locus *l*, and seek to probabilistically assign individuals to each of these populations given their genotypes. When allowing for admixture, an additional parameter *Q* models the proportion of each individual’s genome that originated from a given population. While other programs differ from structure in using variational inference (`fastStructure;` Raj et al. 2014) or a maximum likelihood framework (`ADMIXTURE;` Alexander et al. 2009; `FRAPPE;` Tang et al. 2005), they are united in proposing an explicit causal model for input data, assuming linkage equilibrium between loci and Hardy Weinberg equilibrium between alleles. In contrast, nonparametric methods such as principal components analysis and *K*-means clustering (Jombart et al. 2010; Novembre et al. 2008) first reduce the dimensionality of an allele frequency matrix and then seek to identify groups of individuals that minimize an objective function without explicitly modelling the attributes of genetic data.

Because of these differences, parametric and nonparametric approaches may show different sensitivities to SFS generated through biased data collection methods. It’s possible these sensitivities also reflect the influence of the type datasets available during each program’s initial development: for example, as structure’s underlying algorithm was tested prior to widespread adoption of high throughput sequencing methods and initially applied on microsatellite data screened for appropriate frequency distributions (Pritchard et al. 2000, Li et al. 2000), the characteristics of unfiltered modern SNP datasets may present unanticipated challenges to accurate population genetic inference. Yet to the best of our knowledge, no studies have directly addressed this potential source of error in population genetic and phylogeographic studies. Here, we assess the influence of minor allele frequency (MAF) thresholds on inference of population structure. We evaluate the ability of model-based and nonparametric clustering methods to describe population structure in both simulated and empirical genomic datasets and find that structure is confounded by singletons and both approaches are sensitive to variation in MAF thresholds. We propose a simple hypothesis to explain this behavior and recommend a set of best practices for researchers seeking to describe population `structure` using reduced-representation libraries.

## Methods

### Simulated Data

We simulated genome-wide SNP datasets under a custom demographic model in `fastsimcoal2` (Excoffier et al. 2013) in order to assess the impacts of MAF filtering on population structure inference in the absence of sequencing or assembly error. Model parameters were chosen to reflect a plausible demographic history for our empirical case (see below), with one population experiencing successive splits 60,000 and 40,000 generations in the past after which all populations increase in size exponentially, reaching a final *N_e_* of 50,000 for the “outgroup” lineage and 500,000 for the remaining populations (**Figure 1A**). Migration is allowed among all populations after the final divergence event. We included a mutation rate parameter of 2×10^−6^ in simulated data—equivalent to selecting a single SNP from a 200 bp region in an organism with an average genome-wide mutation rate of 1×10^−8^ (see `fastsimcoal2` user manual). Missing data—a common feature of reduced-representation library SNP datasets—was simulated by randomly dropping 25% of the alleles at each simulated locus.

**Figure 1:**
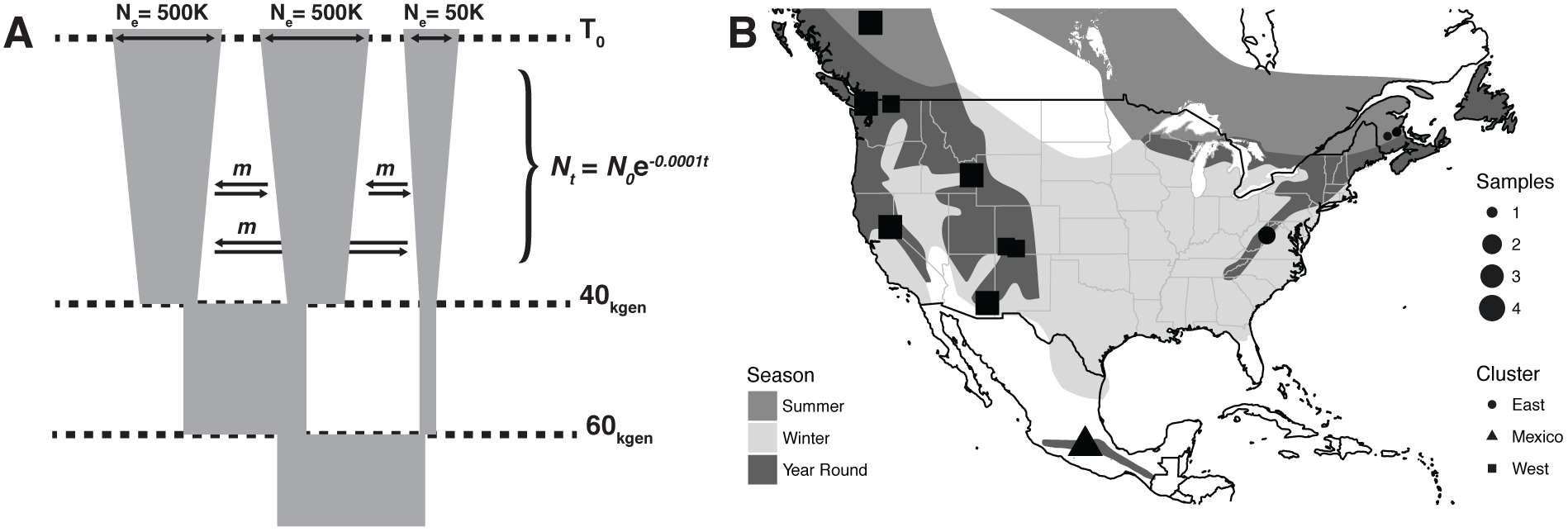
(A) The demographic model used in simulating SNP datasets. (B) Sampling localities and sizes for *Regulus satrapa*, with *a priori* population assignments.

We generated 10 independent simulations using the same starting parameter values and replicated analyses 10 times for each dataset. Each simulation was initialized with 5,000 loci across 10 individuals in each of the 3 populations. After converting `fastsimcoal2` output to `structure’s` input file format, we generated MAF-filtered datasets at each of the following cutoffs: 1/60, 2/60, 3/60, 4/60, 5/60, 8/60, 11/60, and 15/60.

To test whether variation in inferred admixture levels was caused by MAF thresholds specifically rather than a drop in the total size of the data matrix after filtering, we reran the above simulations but initialized with 40,000 loci and then randomly downsampled all alignments to 1000 sites after applying MAF cutoffs.

### Empirical Data

We collected genome-wide SNP data from 40 individuals of the widespread North American passerine *Regulus satrapa*, the Golden Crowned Kinglet. Our geographic sampling aimed to represent three areas of the species’ breeding range a previous study with mitochondrial DNA suggested were distinct populations (Klicka 2017, unpublished data): subspecies *satrapa* in the Eastern US / Canada; subspecies *olivaceous* / *apache* in the coastal and Rocky Mountain US / Canada, respectively; and subspecies *azteca* in the Sierra Madre del Sur and Transvolcanic Belt of Mexico (**Figure 1B**). We extracted whole genomic DNA using Qiagen DNEasy extraction kits and prepared reduced-representation libraries via the ddRADseq protocol (Peterson et al. 2012) using the digestion enzymes Sbf1 and Msp1 and a size-selection window of 415-515 bp. We sequenced the resulting libraries for 50 bp single-end reads on an Illumina HiSeq 2500.

We assembled reads into sequence alignments *de novo* using the program `ipyrad` v. 0.7.11 (https://github.com/dereneaton/ipyrad). We set a similarity threshold of 0.88 for clustering reads within and between individuals, a minimum coverage depth of 6 per individual, and a maximum depth of 10,000. To exclude paralogs from the final dataset, we filtered out loci sharing a heterozygous site in 50% of samples. We define “locus” throughout this manuscript as a cluster of sequence reads putatively representing the same 50 bp region downstream of an Sbf1 cut site. Because missing data can have a strong influence on population genetic inference (Arnold et al. 2013, Gautier et al. 2013) and preliminary exploration suggested anomalous clustering behavior, we removed 7 individuals from our dataset prior to all downstream analysis. Of these final 33 samples, we required each locus to be sequenced in at least half of samples and randomly selected one SNP per locus.

### Population Structure Analyses

We ran 10 replicate `structure` analyses for all MAF filters of simulated (n=80) and empirical data (n=8) using the correlated allele frequency model with admixture for 250,000 generations each, with 10,000 generations of burn-in. All runs were initialized using a random seed value drawn from a uniform distribution with range (0 – 10,000). No prior population assignment information was included in the model. All other settings were left at default values.

Principal components analysis, *K*-means clustering and discriminant analysis of principal components (DAPC) were conducted using the R package `adegenet` (Jombart et al. 2010) and the MAF-filtered structure files as input. Missing data was replaced with the mean values across the full sample before running PCAs. DAPCs were initialized using the *K*-means clustering solution and tested by training the model on half the individuals in each population, then predicting the population assignment of the remaining individuals. PCA and *K*-means analyses were repeated 10 times per input dataset, and DAPC cross validations were repeated 10 times per *K*-means replicate.

In practice most clustering solutions are assessed visually by comparing bar plots of `structure’s` output or scatter plots of PCs 1 and 2. In order to quantitatively compare clustering results across methods and MAF cutoffs, we estimated two summary statistics: the proportion of correct population assignments, and the ratio of distances between individuals within populations to those between all individuals (we refer to this as “*P C_st_*” to emphasize its analogy to *F_st_* and *ϕ_st_*). The proportion of correct population assignments was estimated by assigning each individual to a single cluster (for `structure` results individuals were assigned to the cluster with the highest *q* value), swapping cluster labels to account for stochastic label switching during inference, and comparing inferred and true population assignments.

Within-to-total population distance ratios were calculated as:

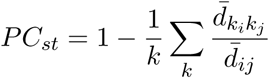

where *k* is the population index, *i* and *j* are the indices of individuals, and *d̄* is the mean Euclidean distance between individuals in a *k*-dimensional space described by the first *k* principal components or the columns of the *q* matrix returned by `structure`. More simply, this ratio is the average distance between individuals in the same population over the average distance between all individuals. High values indicate that inferred clusters are discrete, while low values indicate that clusters overlap—either reflecting uncertainty in individual assignments or admixture among populations.

## Results

### Simulations and sequence assembly

Following MAF filtering, our simulated datasets retained an average range of 3942 (for MAF=1) to 242 (for MAF=15) loci. Constant-length datasets were always subsampled to 1000 bp. For our *Regulus satrapa* ddRAD libraries, Illumina sequencing returned an average of 781,011 quality-filtered reads per sample. Clustering within individuals identified 35,722 putative loci per sample, with an average depth of coverage of 22x. After clustering across individuals and applying paralog and depth-of-coverage filters, we retained an average of 4286 loci per sample. Prior to applying MAF filters and removing individuals for excess missing data, our alignment included 3898 unlinked diallelic SNPs from sequenced in at least 30 of the original 40 samples. Our final MAF-filtered datasets ranged from 3419 (MAF=1) – 431 (MAF=20) loci.

### Parametric clustering

The ability to detect population subdivision in both simulated and empirical datasets varied widely across MAF thresholds using the model-based method `structure` (**Figure 2**). In both constant and variable-length datasets, including singletons caused structure to assign all individuals to the same majority ancestry cluster. For variable-length simulated datasets, after excluding alignments with singletons, higher MAF thresholds are also associated with lower population discrimination (linear regression, *p*<2e-16, *R*^2^=0.21, slope= −0.02, *df* =698) and assignment accuracy (*p*<2e-16, *R*^2^=0.20, slope=-0.014, *df* =698) (**Figure 5**). The association between high MAF cutoffs and population discrimination is reversed in constant-length datasets – more stringently filtered datasets infer more discrete clusters—though the effect is much weaker (*p*<1e-10, *R*^2^=0.093, slope = 0.007).

**Figure 2:**
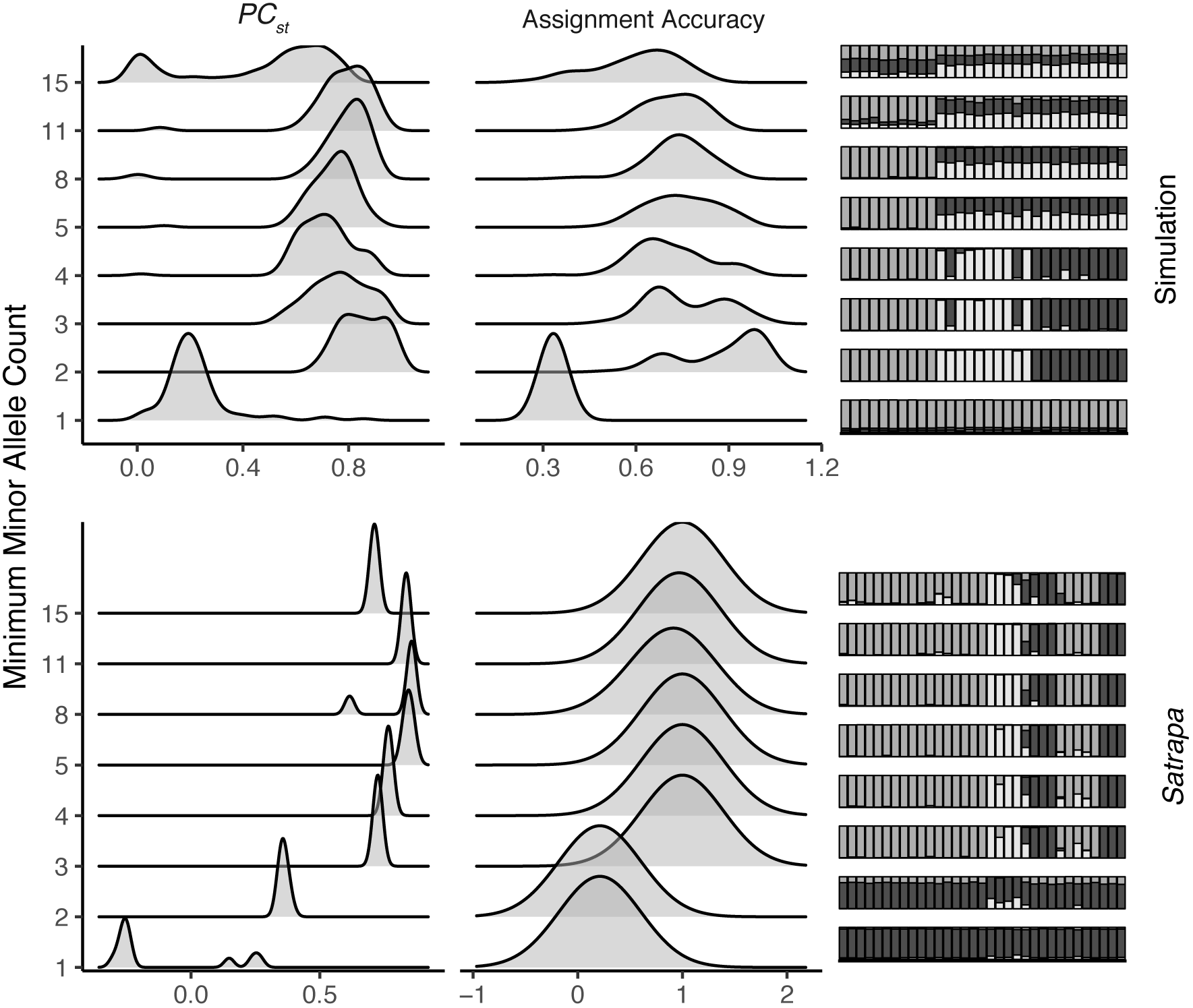
The influence of minor allele count on structure’s assignment accuracy and *P C_st_* for simulated and empirical datasets

### Nonparametric clustering

In contrast to `structure`, both *K*-means clustering accuracy and *P C_st_* were robust to inclusion of singletons (**Figure 3**). However, both measures were highly sensitive to MAF thresholds in simulated data. Both *P C_st_* and *K*-means assignment accuracy decline as the MAF threshold is increased (linear regression, *P C_st_*: *p*<2e-16, *R*^2^=0.606, slope=-0.01; *K*-means accuracy: *p*<2e-16, *R*^2^=0.404, slope=-0.011, *df* =798). As with structure these relationships are reversed but weaker when alignment length is held constant (linear regression, *P C_st_*: *p*<2e-16, *R*^2^=0.216, slope=0.005; *K*-means accuracy: *p*<2e-16, *R*^2^=0.069, slope=0.006) (**Figure 6**), though the relationship remains negative across MAF cutoffs in the range of 0.01 0 – 0.083. For empirical data, both methods achieved near-perfect assignment accuracy under all MAF cutoffs.

**Figure 3:**
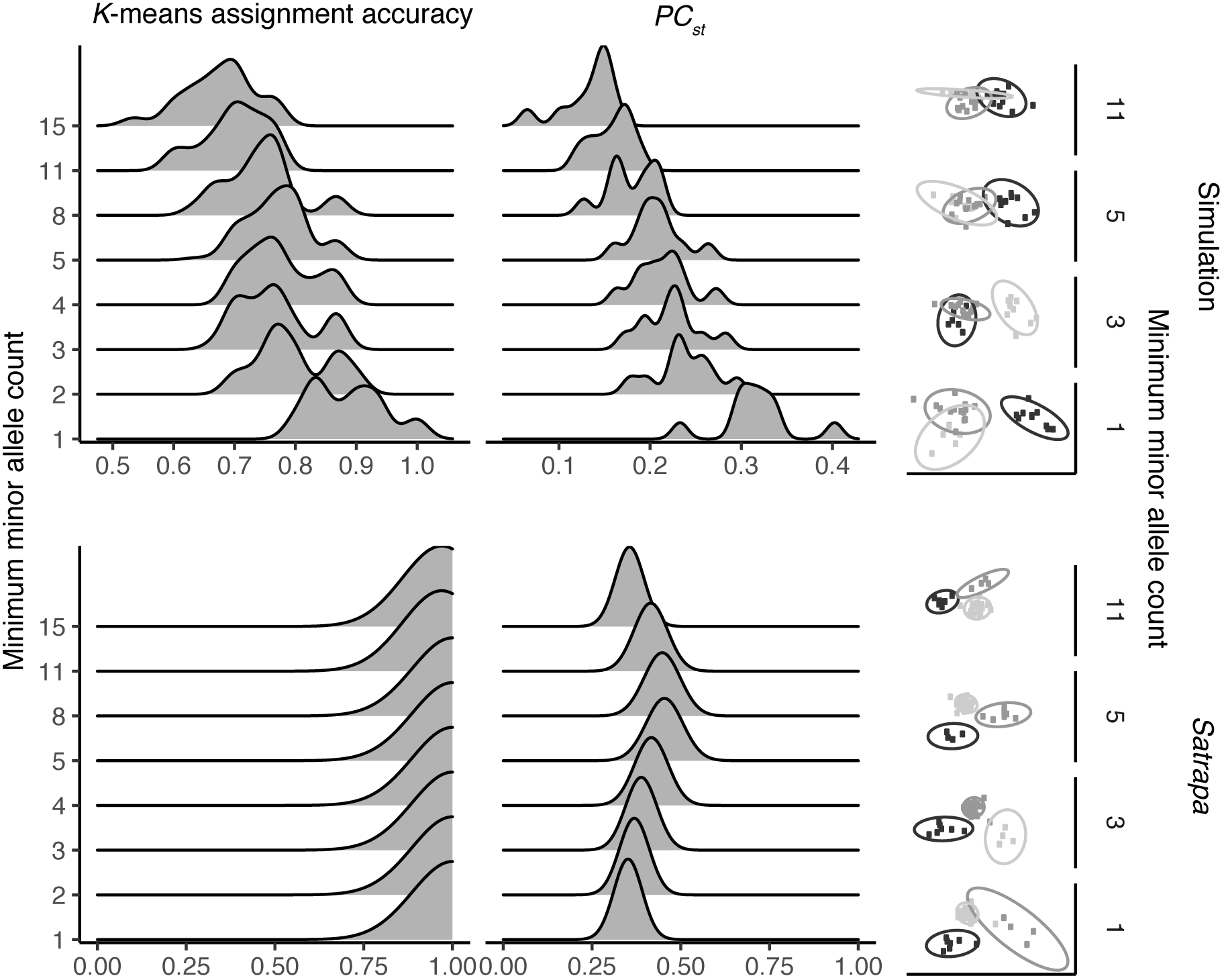
The influence of minor allele count on *K*-means assignment accuracy and *P C_st_* for simulated and empirical datasets

## Discussion

### Inference of population structure is sensitive to MAF

Our results demonstrate that inference of population structure can be strongly influenced by choice of MAF threshold with both model-based and multivariate approaches. `Structure` fails to detect even moderate population subdivision (*F_st_ ≈* 0.05) when singletons are included in the alignment, and both methods generally infer increasing levels of admixture as the minimum MAF of sites included in the alignment is increased. These trends do not occur when alignment length is held constant, suggesting that most of the effect is driven by a drop in the total size of the data matrix after filtering by MAF. In practice this will occur in most empirical datasets when genotypes are estimated from sequencing data. For chip-based approaches in which SNPs are first screened for variation at some cutoff, our analysis suggests that clustering results should be relatively robust to implicit MAF cutoffs applied during chip design.

Two factors may explain the pattern of increased admixture in more stringently filtered datasets: variation in the total size of the data matrix, and the distribution of mutations on a coalescent tree. In simulated datasets with varying size (as in nearly all empirical cases), increasing the MAF cutoff decreases the total size of the data matrix and leads to much higher estimates of individual admixture. This is in part an interpretive issue, as the strong effect of the size of the data matrix suggests that the high *q*-matrix values reflect uncertainty in individual assignments rather than higher admixture levels. However, because parametric approaches are typically interpreted in light of their generative model, many users are likely to see this pattern as evidence of higher gene flow.

A secondary cause of increased admixture in more stringently filtered datasets is the time distribution of mutations in a coalescent tree. Under a standard coalescent model the expected number of sites with a derived allele present in *i* samples (*s_i_*) is the total length of branches subtending *i* descendents (*τ_i_*), multiplied by the expected number of mutations per unit time (*θ/*2):

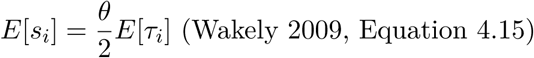

Low-frequency alleles represent mutations that occurred on branches with few descendants, and these branches are typically found close to the present (**Figure 4**; see Appendix 1 for simulation details). Low-frequency alleles therefore contain a disproportionate amount of information about recent events. Removing them is similar to drawing a horizontal line across a coalescent tree and dropping mutations that occur beyond that line. In the absence of recent pulses of gene flow, this causes populations to appear less differentiated as the MAF threshold increases, seen in PCA output as reduced distance between clusters and in structure output as increased admixture within individuals (though some would argue that this is simply a misinterpretation of `structure’s` output, e.g. Lawson et al. 2017).

**Figure 4:**
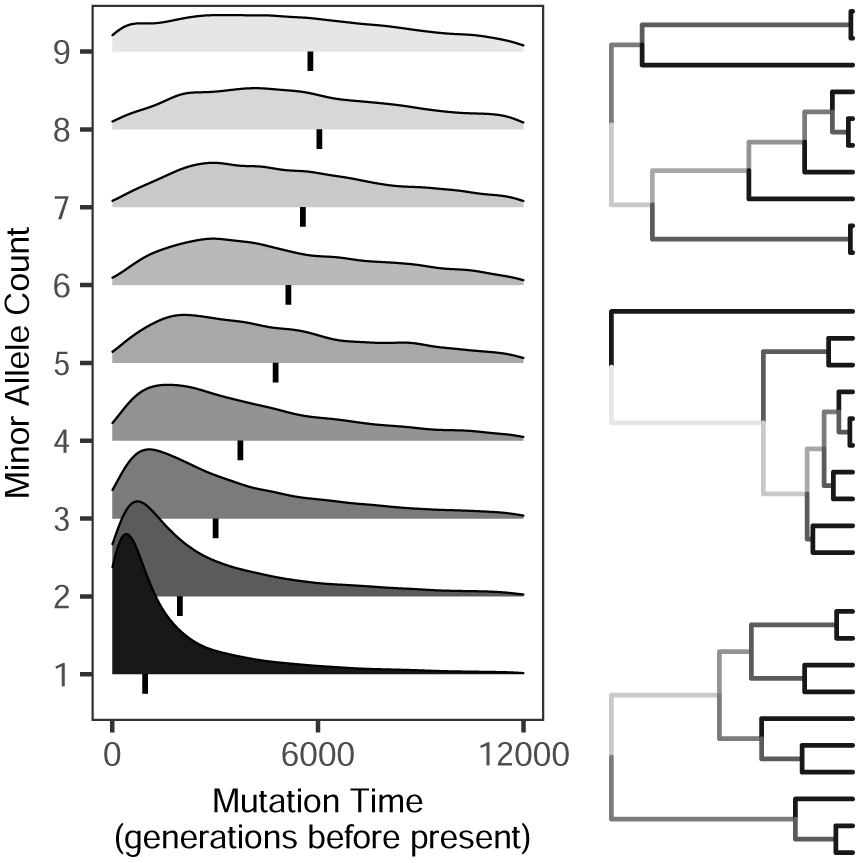
Time distribution of mutations with varying derived allele counts

Model-based analyses’ failure to recover a clear signal of population subdivision when singletons are included in the alignment is more difficult to explain. The issue appears to be related to overfitting as a result of either a high frequency of uninformative rare alleles or a high frequency of uninformative common alleles (Alexander and Lange 2011). In both scenarios, population *k*_1_ receives an allele frequency distribution averaging out true population specific-frequencies of common alleles, resulting in the broad band of majority ancestry visible in **Figure 2**. Subsequently, populations *k*_2_… *k_n_* receive high frequencies of singletons or otherwise uninformative rare alleles, resulting in the additional bands of minority ancestry shared across all individuals. With our simulated data, rare but non-singleton alleles reflect fine population structure and thus harm inference when excluded; with our empirical data, rare alleles are uninformative and serve only as noise to common allele frequency distributions reflecting true population history.

This hypothesis is consistent with a pathology related `structure’s` inability to model mutation of modern alleles, previously identified as a potential obstacle to accurate inference of population structure under certain histories (Shringapure and Xing 2009). Because structure assumes each unique allele in the input dataset has a distinct frequency in its parent population, recent mutations— e.g., derived alleles—are erroneously treated as representative of a separate population-specific allele frequency profile rather than as descendants of ancestral copies. If a sufficient number of singletons are present in the dataset, the noise from these false allele frequency profiles may mask the signal from alleles indicative of “true” populations. Though most multivariate analyses were robust to inclusion of singletons, a similar pattern of low accuracy and population discrimination was observed in PCA when alignment length was held constant—likely because low-frequency alleles hold less information about inter-group differences than moderate-frequency alleles, and low-frequency alleles will be a larger proportion of the total data matrix in this case.

### Recommendations for setting MAF thresholds in population genetic studies

Our results suggest that SFS distributions that can cause structure and other model-based programs to erroneously fail to detect structure may be generated by either normal demographic processes (e.g., exponential population growth with relatively recent divergence, as in our simulated example) or by assembly errors (potentially present in our empirical example, and well documented in other de novo RADseq datasets, e.g. Shafer et al. 2016). As a consequence, a broad set of empirical studies may be affected. We recommend researchers using model-based programs to describe population structure observe the following best practices: 1) duplicate analyses with nonparametric methods such as PCA and DAPC with cross validation; 2) exclude singletons; and 3) compare alignments with multiple assembly parameters. When seeking to exclude only singletons in alignments with missing data (a ubiquitous problem for reduced-representation library preparation methods), it is preferable to filter by the count (rather than frequency) of the minor allele, because variation in the amount of missing data across an alignment will cause a static frequency cutoff to remove different SFS classes at different sites. The scripts used to filter structure input files for this manuscript are available at https://github.com/cjbattey/LinckBattey2017_MAF_clustering.

### *Population genetics of* Regulus satrapa

Though describing population structure and phylogeographic patterns of the Golden-crowned Kinglet was not the primarily goal of our study and will be elaborated on elsewhere, our data provide novel evidence for deep splits across the range of the species, corroborating previous mtDNA evidence (Klicka 2017, unpublished data). Curiously, the results of our model-based population structure inference suggest not only singletons but also doubletons have a high noise to signal ratio, while common alleles (MAC ≥ 3) accurately reflect expected relationships. This pattern may be driven by either purifying selection eliminating geographically localized variants (Nelson et al. 2012, Jackson et al. 2015), a population bottleneck (Nei et al. 1975; Gattepaille et al. 2013), a burst of recent migration following exponential population growth (Slatkin 1985), or assembly artifacts resulting in a high proportion of uninformative / erroneous sites (Shafer at al. 2016). While all scenarios are likely contributing to some extent, studies of genetic variation in similar taxa provide support for post-Pleistocene expansion and gene flow among populations separated by ice sheets (Spellman and Kilcka 2006), processes that may result in similar SFS distributions to our example.

### Future directions

With simulated and empirical cases reflecting similar (if non-identical) site frequency spectra, our focus was on a necessarily narrow range of demographic scenarios and a relatively narrow range of SFS distributions. Future examinations of the sensitivity of population genetic inference to MAF thresholds with datasets simulated under a diversity of evolutionary histories may shed light on the biological processes generating problematic SFS, and lead to the development of more robust model-based programs. While other parametric population structure inference programs share `structure’s` underlying model and we believe the broad patterns reported here will be similarly reflected, differences in implementation (e.g., MCMC mixing) may shape specific sensitivities. A broader survey of model-based population structure inference methods will help clarify which approaches are best suited to NGS data, and lead to the development of more robust software for describing the fundamental units of biological organization.

## Acknowledgements

This research was supported by a National Defense Science and Engineering Graduate (NDSEG) Fellowship to E.B.L. Comments on a preprint version of this article significantly improved its contents. We especially thank Dave Slager for his early contributions to our thinking on this problem.

## Appendix 1: Coalescent simulations of mutation timing

To demonstrate the relationship between the time a mutation occurs and the minor allele frequency at a given site, we developed a single-locus discrete coalescent simulator that tracks the generation of each mutation. Simulations are initialized with a set of individuals at generation 0. At each generation moving into the past, the probability of a coalescence event is approximated by sampling from a binomial distribution with probability of success (*k –* 1)*/N*, where *k* is the number of tracked lineages in the current generation and *N* is the total population size. If a coalescent event occurs, two individuals are randomly selected and merged. The number of mutations occurring on the branches joining the coalescing individuals is then approximated by conducting *n* binomial draws with probability *μ*, where *n* is the branch length and *μ* is the mutation rate. Mutations are then assigned to generations by randomly sampling values across the range of generations between successive coalescent events. Results are stored in a Newick tree and a set of tables, and plotted with functions from the R packages ape (Paradis et al. 2004), ggplot2 (Wickham 2016), and ggridges (Wilke 2018). The simulator is implemented in R and is available at: https://github.com/cjbattey/coalesceR.

## Supplemental Material

**Figure 5:**
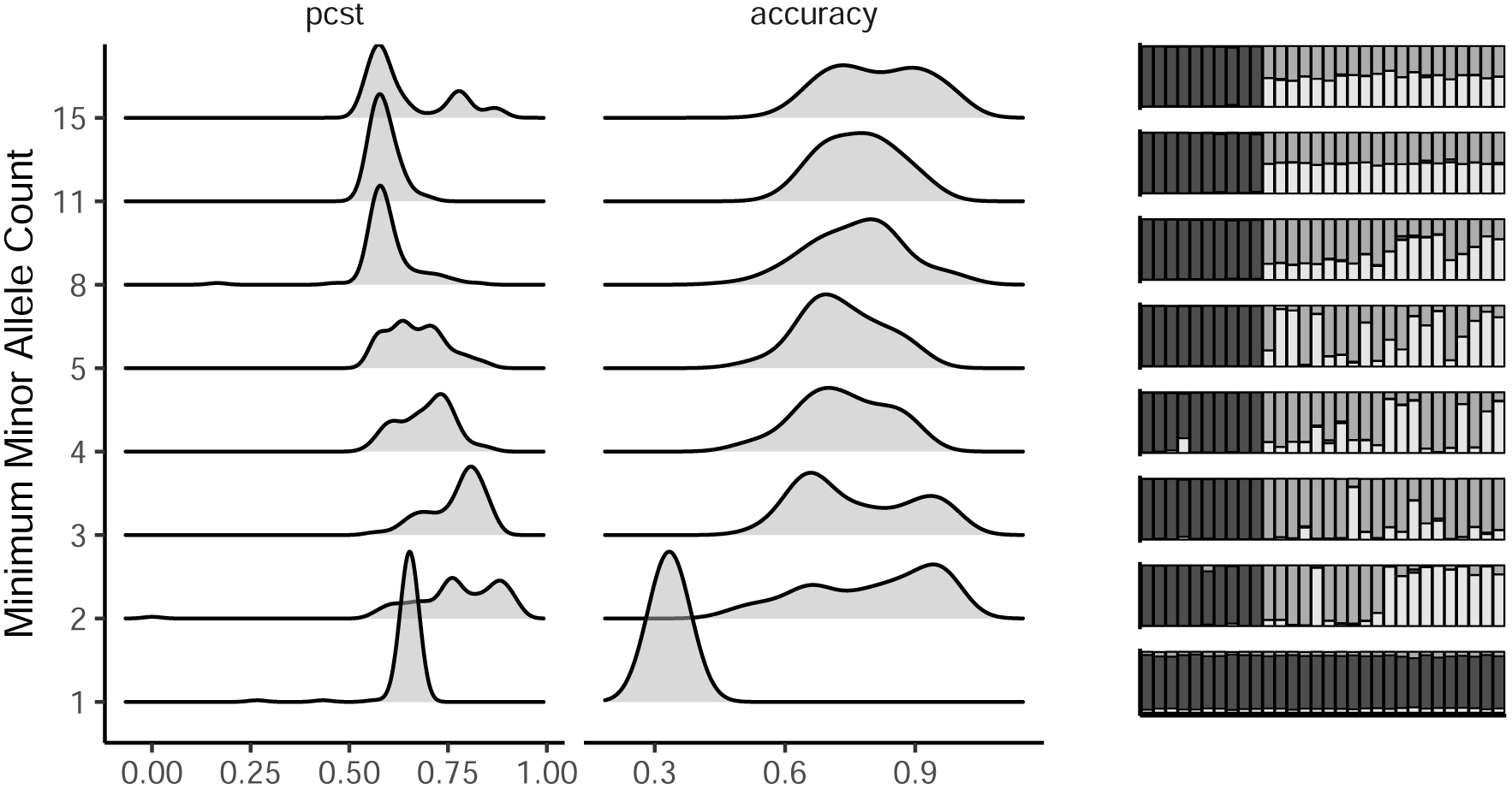
`Structure` results for simulated data when alignment length is held constant at 1000 SNPs.

**Figure 6:**
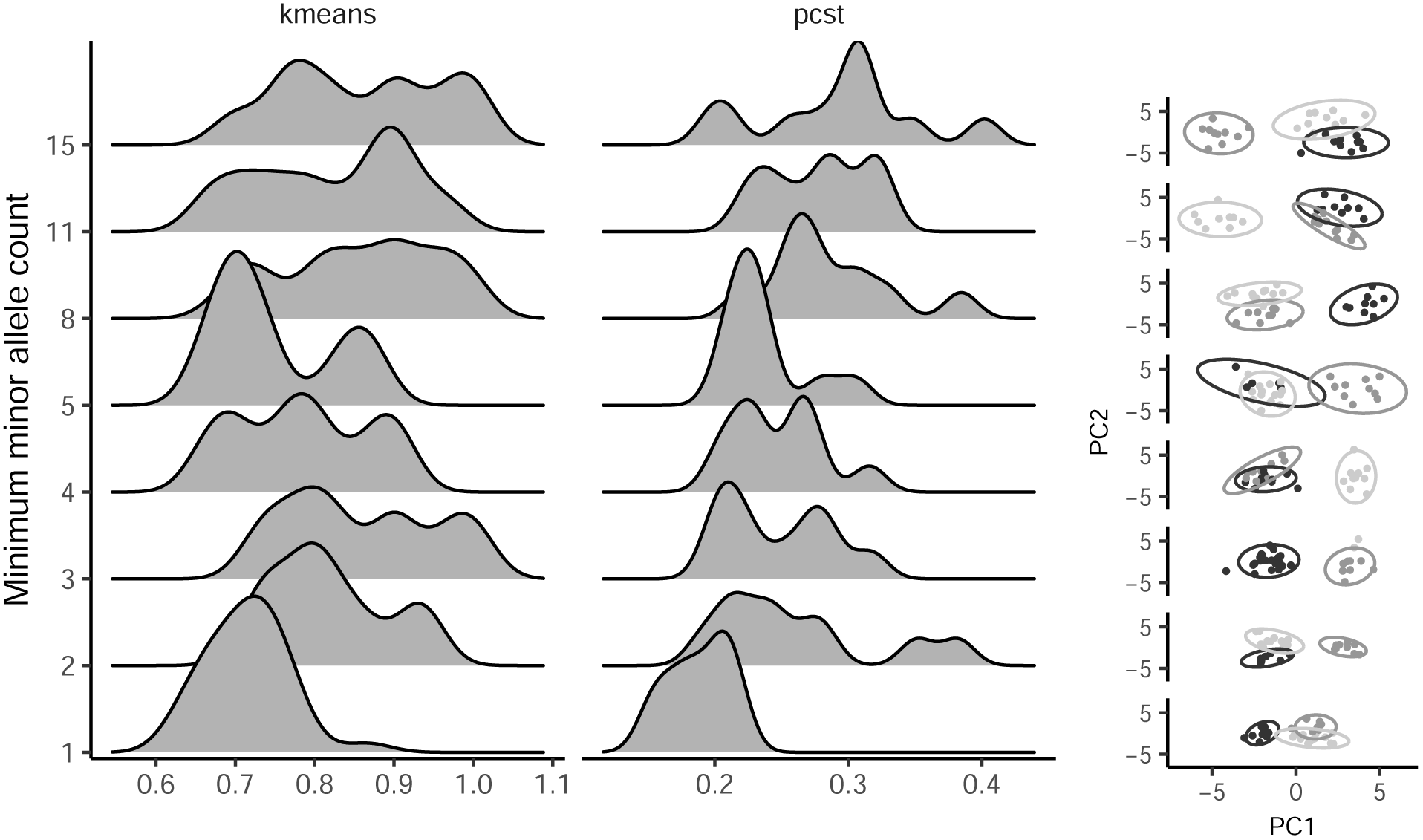
Multivariate clustering of simulated data when alignment length is held constant at 1000 SNPs.

